# Growing old too early, automated assessment of skeletal muscle single fiber biomechanics in ageing R349P desmin knock-in mice using the *MyoRobot* technology

**DOI:** 10.1101/737973

**Authors:** Charlotte Meyer, Michael Haug, Barbara Reischl, Gerhard Prölß, Thorsten Pöschel, Stefan J Rupitsch, Christoph S Clemen, Rolf Schröder, Oliver Friedrich

**Author notes:** **For correspondence:** (FMS). Authors contributed equally to this work (Charlotte Meyer, Michael Haug).

## Abstract

Muscle biomechanics is determined by active motor-protein assembly and passive strain transmission through cytoskeletal structures. The extrasarcomeric desmin filament network aligns myofibrils at the z-discs, provides nuclear-sarcolemmal anchorage and may also serve as memory for muscle repositioning following large strains. Our previous analyses of R349P desmin knock-in mice, an animal model for the human R350P desminopathy, already depicted pre-clinical changes in myofibrillar arrangement and increased fiber bundle stiffness compatible with a pre-aged phenotype in the disease. Since the specific effect of R349P desmin on axial biomechanics in fully differentiated muscle fibers is unknown, we used our automated *MyoRobot* biomechatronics platform to compare passive and active biomechanics in single fibers derived from fast- and slow-twitch muscles from adult to senile mice hetero- or homozygous for this desmin mutation with wild-type littermates. Experimental protocols involved caffeine-induced Ca^2+^-mediated force transients, pCa-force curves, resting length-tension curves, visco-elasticity and ‘slack-tests’. We demonstrate that the presence of R349P desmin predominantly increased single fiber axial stiffness in both muscle types with a pre-aged phenotype over wild-type fibers. Axial viscosity was unaffected. Likewise, no systematic changes in Ca^2+^-mediated force properties were found. Notably, mutant single fibers showed faster unloaded shortening over wild-type fibers. Effects of ageing seen in the wild-type always appeared earlier in the mutant desmin fibers. Impaired R349P desmin muscle biomechanics is clearly an effect of a compromised intermediate filament network rather than secondary to fibrosis.

## Introduction

Skeletal muscle is the largest organ system of the body and under constant mechanical stress, either due to passive strain or through active contraction producing axial and lateral stresses. While lateral forces are distributed between single fibers across anchorage points in the extracellular matrix (ECM) to the intracellular cytoskeleton via the dystrophin-glycoprotein complex (DGC) ***Ramaswamy et al. (2011)*** and focal adhesion complexes ***Ra et al. (1999)***, axial forces are distributed through contractile (active) and non-contractile (passive) elements. Apart from the giant, roughly 1.5*μ*m long elastomeric protein titin, being responsible for the visco-elastic properties of single muscle fibers through unfolding of globular domains under strain ***Mártonfalvi and Kellermayer (2014); Powers et al. (2018)***, connecting proteins of the extra-sarcomeric intermediate filament (IF) family may also be a vital determinant of axial elasticity. An important member of the IFs is the type III filament protein desmin, transversely linking adjacent myofibrils at the level of the z-disc and thus, being responsible for myofibrillar register ***Waterman-Storer (1991)***; ***Anderson et al. (2001); Meyer et al. (2010)***. In humans, desmin is encoded on chromosome 2q35 by a single copy gene. The 53 kDa desmin presents a tripartite structure with a central-helical coiled-coil domain flanked by non-helical tail and head domains. Due to its intrinsic self-assembling properties, it builds three-dimensional networks, starting with supercoil formation via dimerization of two desmin molecules. Two such dimers then associate into tetramers that represent the repetitive add-on units for spontaneous assembly of 60 nm long filaments, the so-called unit-length filaments (ULFs, ***Clemen et al. (2013))***. Serial longitudinal annealing of ULFs consequently builds short filaments, extending the IF network. In the end, long filaments reduce their diameter by spontaneous radial compaction to form the mature IF network. The network connects to multiple intracellular adhesion sites by cross-bridging proteins from the spectrin superfamily, i.e. plectin and nesprins ***Liem (2016)***. In skeletal muscle, IFs form a huge stress-transmitting and stress-signalling network, and desmin in particular, is required for the maintenance of myofibrillar alignment, nuclear positioning and shape, stress production and sensing ***Palmisano et al. (2015); Meyer et al. (2010)***. Due to the low turn-over rates of IF proteins, the IF network remains largely intact even when exposed to large physical strains, e.g. surviving at least 350% strains before rupture ***Block et al. (2017); Kreplak et al. (2008)***. This led to their proposed role of acting as a cytoskeletal ‘position-memory’ to ensure the proper re-assembly of cytoskeletal components following recovery from large strains ***Gan et al. (2016)***. The deleterious effects of abnormal desmin IF networks, due to either the additional presence of mutant or the complete lack of wild-type desmin protein, are emphasized by the group of human desminopathies that comprise autosomal-dominantly and recessively inherited myopathies and cardiomyopathies ***Clemen et al. (2013)***. Human desminopathies are clinically characterized by a broad phenotypic variability ranging from primary distal myopathies, limb girdle muscular dystrophies and scapuloperoneal syndromes to generalized myopathies ***Walter et al. (2007); Baer (2005); Clemen et al. (2009); Durmuş et al. (2016)***. The major problem with elucidating pathophysiological mechanisms of the human phenotypes is that knowledge about early and intermediate stages of the disease is usually elusive, since muscle tissue specimens for research are not available from patients at pre-clinical stages. Therefore, a patient-mimicking knock-in mouse strain carrying the R349P desmin mutation, the murine orthologous of the human R350P mutation, was generated ***Clemen et al. (2015)***. This model already allowed detailed systematic studies of clinical and myopathological phenotypes as well as age-dependent effects on the disease progression in heterozygous (het) and homozygous (hom) desminopathy mice over their wild-type (wt) littermates ***Diermeier et al. (2017a)***. In particular, our previous work demonstrated that the expression of R349P mutated desmin compromised the three-dimensional arrangement and the order of the myofibrillar lattice already starting in young mice before presenting muscle. The latter findings were interpreted as a pre-aged phenotype of muscle structural ageing in the R349P environment ***Diermeier et al. (2017b)***. Moreover, biomechanical analyses of small fiber bundles, initially in slow-twitch, load-bearing *M. soleus* (SOL) from young het and hom R349P desmin mice, showed a marked increase in passive bundle stiffness compared to wt bundles ***Clemen et al. (2015); Diermeier et al. (2017a)***. Since robust biomechanical experiments in small muscle fiber bundle preparations are difficult to carry out and require precise actuation to record steady-state resting length tension curves at slow strain speeds, we engineered a novel automated biomechatronics system that also contains a sensitive force transducer technology for recordings of active and passive axial muscle forces ***Haug et al. (2018)***. Alongside with the ongoing engineering progress, our system was also systematically validated extending our previous recordings in SOL bundles from R349P desmin mice not only to the fast-twitch extensor digitorum longus (EDL) muscle, but also to a wide age range from young (17-23 weeks) to aged (60-80 weeks) animals ***Haug et al. (2019)***. Again, our findings in young animals of increased tissue stiffness were confirmed in both muscle entities with a pre-aged phenotype in the desminopathy model. However, since also ECM re-modeling has been shown with increased levels of tissue fibrosis with age in the R349P background ***Diermeier et al. (2017a)***, an unambiguous explanation towards the link of increased axial stiffness to the disrupted desmin network could not be drawn. This is because in small fiber bundles, both the ECM and the intracellular cytoskeleton still contribute to the overall axial compliance. To tackle this constraint, the present study was designed to revisit biomechanical tests in EDL and SOL R349P desminopathy muscles using pure mechanically dissected single fiber segments with a refined version of our *MyoRobot* suitable to record single fiber forces. Our results provide novel insights into (i) the connection of mutated desmin to axial active/passive biomechanics in single fibers and (ii) the age-dependent progression of altered fiber mechanics in this desminopathy model.

## Results

### Ca^2+^-mediated active isometric force and contractile apparatus Ca^2+^-sensitivity automatically assessed in single fibers from R349P desminopathy SOL and EDL muscles during ageing

Fig. 1A shows representative *MyoRobot*-executed force transient recordings of a caffeine-triggered Ca^2+^-mediated force response to empty the SR of its releasable Ca^2+^, followed by a maximum Ca^2+^-saturated activation of the contractile apparatus in HA solution. Finally, exposure to EGTA-rich HR solution buffered any remaining excess Ca^2+^ ions. Consistent with the characteristics of fast-vs. slow-twitch muscle, EDL and SOL fibers showed faster or slower transient kinetics, respectively. Caffeine-induced peak force (Fig. 1B), maximum Ca^2+^-saturated force amplitudes (Fig. 1C) and their ratio (Fig. 1D) were evaluated for all single fibers from all genotypes over all ages in both muscles. In EDL, caffeine-induced force developed differentially with age in the three genotypes. In wt single fibers, force amplitudes initially increased with age to significantly drop again in senile animals. In contrast, in the R349P desmin knock-in background, force developed oppositely in het fibers (decrease in the aged group and recovery to adult levels in the senile group) or did not vary significantly within the hom group. Within age groups, we discovered isolated, genotype-specific significant differences that were however, not systematic (Fig. 1B). Unlike caffeine-induced force, maximum Ca^2+^-saturated force was unchanged in EDL single fibers, regardless of age or genotype (Fig. 1C). The ratio of caffeine-induced to maximum force amplitudes serves as an indicator of SR Ca^2+^ filling and thus, showed a similar behaviour as the former (Fig. 1D). While maximum force amplitudes in SOL single fibers were generally similar to those in EDL fibers (Fig. 1C), caffeine-induced force levels were roughly two-times smaller (Fig. 1B). Within SOL fibers, no difference among genotypes was seen while age had a strong negative effect on force amplitudes, which were significantly reduced in wt preparations for all progressing ages, and in het/hom fibers between the adult and the senile age group. Maximum attainable force levels were also impeded by age and displayed a significant decline during ageing within each genotype. Particularly hom fibers were already significantly reduced in the adult age cohort, while the still better performing wt and het fibers gradually declined to the level of hom fibers with age. This suggests a pre-aged phenotype in hom fibers regarding maximum contractile forces. The combined differences regarding force ratios were restricted to a significant age-related, genotype-specific decline (Fig. 1D).

**Figure 1.**
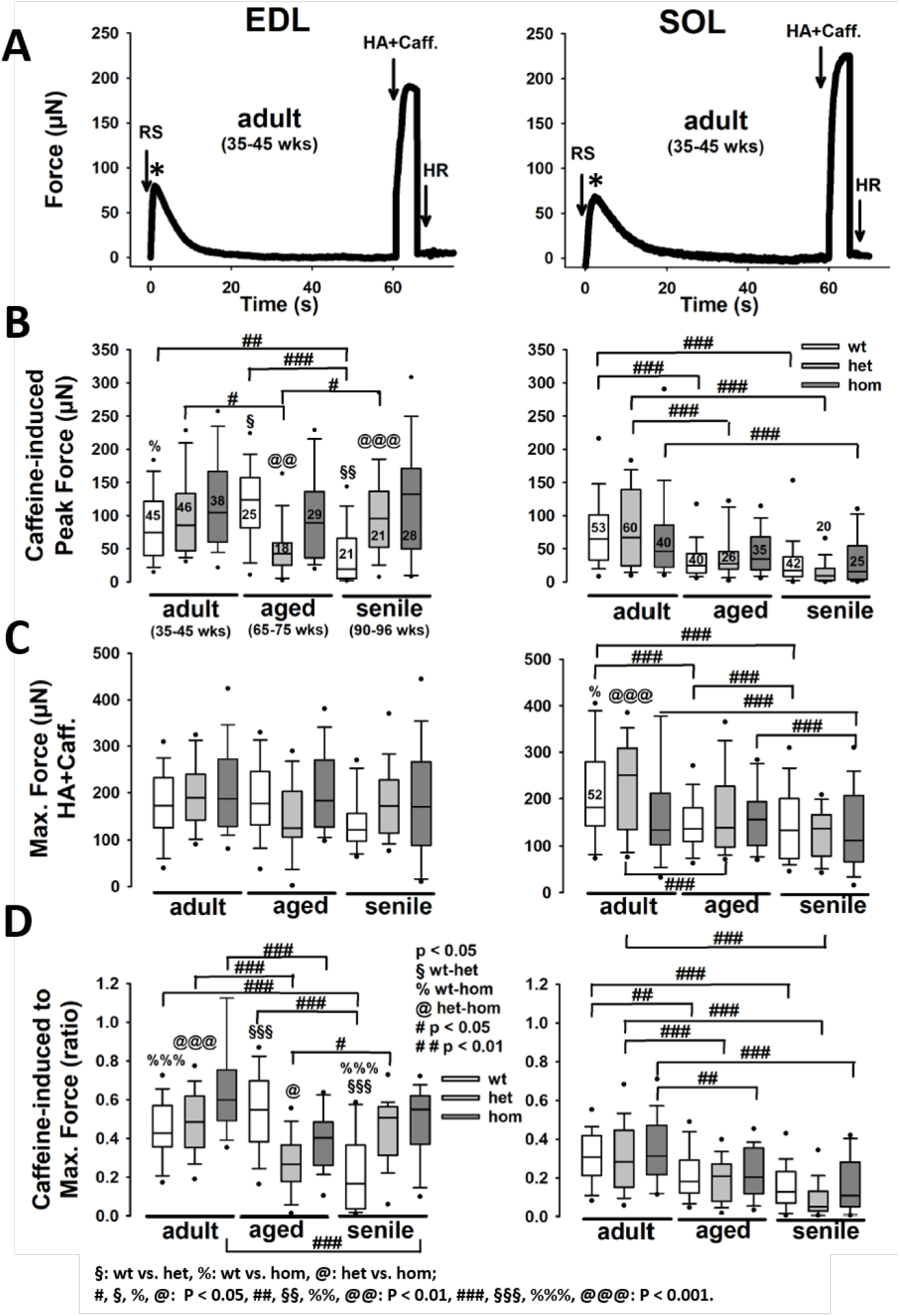
Caffeine-induced force and maximum Ca^2+^-saturated force recorded in permeabilized single EDL and SOL fibers from adult, aged and senile R349P desmin mice. **A**, representative force recordings in a single EDL (left) and SOL (right) fiber. Group analysis of peak force amplitude during caffeine-release (RS) (**B**), steady-state maximum force (HA) (**C**) and respective RS:HAforce ratios (**D**) indicate an overall decrease in SR Ca^2+^-release force during ageing in EDL and SOL, regardless of genotype. Within age groups, RS peakforcewas significantly larger in hom EDL fibers for the adult and senile groups, while they were similar in SOL. In EDL, there was no difference in maximum attainable force among genotypes regardless of age. Thus, RS:HA force ratios in EDL reflect the pattern-differences of RS peaks, while in SOL fibres, relative force during SR Ca^2+^ release over maximum Ca^2+^-saturated force were similar among genotypes and showed a significant decrease with ageing. Significance tested with two-way ANOVAfollowed by post-hoc analysis (Bonferroni). Numbers in box plots: number of single fibers analysed; also valid for (**C**) & (**D**).

To elaborate on the Ca^2+^-sensitivity of the contractile apparatus in single fibers carrying the desmin R349P mutation, pCa-force recordings were performed in single fibers across the three age groups in EDL and SOL muscles, as shown in representative single fiber data traces from each genotype in aged animals (Fig 2A). The top left panels show how force quickly rises to a new steady-state level in response to increasing Ca^2+^ (decreasing pCa) steps. The right panels show the respective average pCa-force of this age group for EDL (left) and SOL (right) muscle along with the average reconstructed Hill fits (Fig. 2B). The curves in Fig. 2A already suggest a marked left-shift of the sensor-curve in R349P desmin knock-in single fibers over wt, indicative of a myofibrillar Ca^2+^ sensitization in presence of mutant desmin. This was also confirmed in the group analysis, where adult hom R349P desmin knock-in EDL single fibers were initially less Ca^2+^ sensitive but became more sensitive than the wt in aged animals. Since in senile mice, an age-related Ca^2+^-sensitization was also observed in wt animals, the behaviour of hom fibers can be considered as a pre-aged phenotype towards higher Ca^2+^-sensitivity of the contractile apparatus. This also agrees with wt single EDL fibers reaching their largest pCa_50_ value one age bin later than hom fibers. Within the oldest age cohort (senile), all pCa_50_ values had finally reached similar levels among genotypes. Unlike EDL, SOL only displayed age-related effects in the wt, with an initial Ca^2+^-desensitization (from the adult to aged animals) that was later revoked in senile animals. Similar to the EDL, adult hom SOL fibers showed yet significantly depressed pCa_50_ values, which however, strongly increased in the aged age cohort while wt fibers only matched those high levels in the senile age group (Fig. 2B). Het fibers showed similar trends as hom fibers, yet, did not reach statistical significance. The Hill coefficients in EDL single fibers showed no significant differences regarding genotypes while age had a significant influence on het fibers between aged and senile animals. In SOL single fibers, differences were present among genotypes, with lower coefficients values for fibers expressing the R349P mutation (except het adult), while age had a significant influence on wt fibers, leading to a significant increase in Hill coefficients in aged and senile fibers over adult fibers, indicative of a higher dynamic range of the myofibrillar Ca^2+^-biosensor complex.

**Figure 2.**
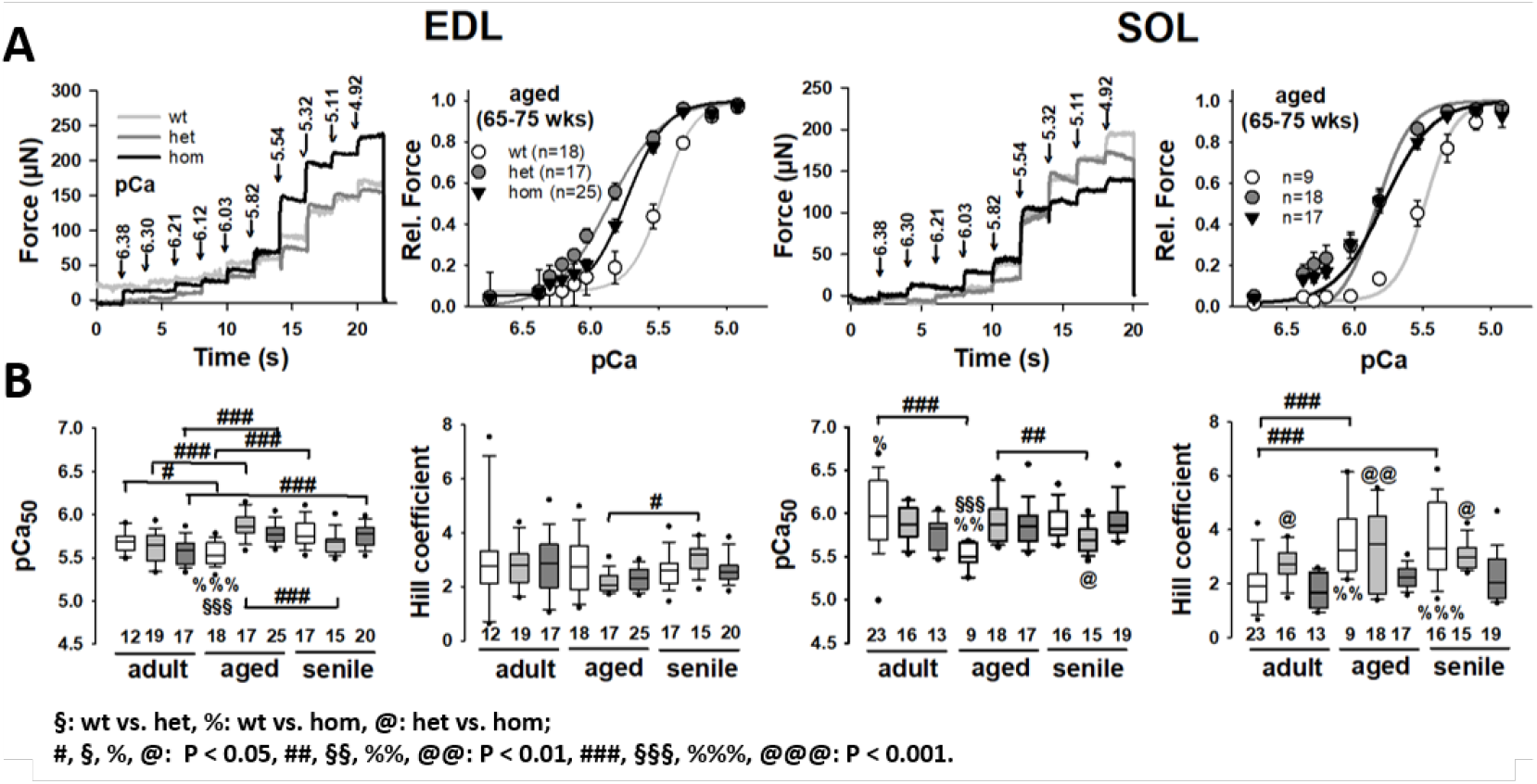
Ca^+^-sensitivity of the contractile apparatus in permeabilized single EDL and SOL fibers from adult, aged and senile R349P desmin mice. **A**, representative force recordings in an aged single EDL (left) and SOL (right) fiber for each genotype showing increasing force for each indicated step change in pCa. The mean pCa-force curves alongside with the mean reconstructed Hill fit to the data are shown to the right. The curves display a marked left-shift in the R349P desmin knock-in background. Group analysis of pCa_50_ values and Hill coefficients in (**B**) show a significantly increased Ca^2+^-sensitivity in aged R349P desmin knock-in animals over the wt which is caught up in the senile group. Likewise, in the adult age group, Ca^2+^-sensitivity is similar between genotypes. In EDL, there was a significant trend towards increasing Ca^2+^-sensitivity in the R349P desmin knock-in background with age, while in SOL, significant age-related changes were only observed in the wt. Overall, differences between wt and hom preparations became more distinct with age. Numbers in box plots: number of single fibers analysed.

### Steady-state resting length-tension curves demonstrate a markedly decreased axial compliance and a pre-aged passive stiffness increase in R349P desmin knock-in single fibers

Our previous work in small fiber bundles from SOL muscles demonstrated an increased axial stiffness in R349P desmin knock-in mice ***Diermeier et al. (2017a); Haug et al. (2019)***. However, since we also documented increased fibrosis in these muscles ***Diermeier et al. (2017a)***, it cannot be ruled out to which extent the observed fibrosis would impact on the increased axial stiffness. To eliminate the influence of ECM components on biomechanics recordings, a preparation of single fibers represented the best possible experimental solution. Fig. 3A shows a series of example RLT curves of single fibers of each genotype and age group from EDL and SOL muscles. The example traces already suggest that the RLT slope strongly increased with age in single fibers with mutation background, the more so in EDL over SOL muscle. This increase occurred in wt EDL single fibers in a less-pronounced fashion, while it was absent in wt SOL samples, which remained at similar levels independent of age. This is in accordance with a pre-aged phenotype in the R349P mutants regarding axial fiber stiffness. As a measure for steady-state stiffness at the end of the stretch to the 140% L_0_ length, the maximum restoration force (FR) was analysed in Fig. 3B, statistically confirming the behaviour seen in the examples. In the adult age group, max. FR values were all similar between genotypes. In the EDL fibers, while they all increased with age, they did so more strongly and earlier in the R349P knock-in background, significantly exceeding the wt in the aged group. By the very old senile age, the wt had then caught up with the mutants. Although not significant, the het fibers had smaller max. FR values than the hom fibers. This trend was also seen in the SOL fibers except for the wt fibers showing significantly decreased max. FR values with age in comparison to the adult group. The higher max. FR values in the R349P knock-in background also impacted on a lower survival of single fibers during stretch. Both, in EDL and SOL muscles, mutant single fibers already broke at lower strains compared to the wt, while fibers het for R349P displayed a better survival than hom fibers (Fig. 3C). Since these results indicate an increased axial stiffness or elasticity, the 10% strain-wise compliance was computed by linear fits to the RLT curve at the indicated strain bins, as described in ***Haug et al. (2019)***, and plotted for all genotypes, ages and strains in Fig.3D. The compliance plots confirm similar mechanical axial compliance for all genotypes in the adult age group while compliance was significantly reduced in mutant fibers of the aged age group. The wt then declined to similar low compliances as the mutants in the senile age group for EDL muscle fibers, whereas for SOL, compliance remained at high levels.

**Figure 3.**
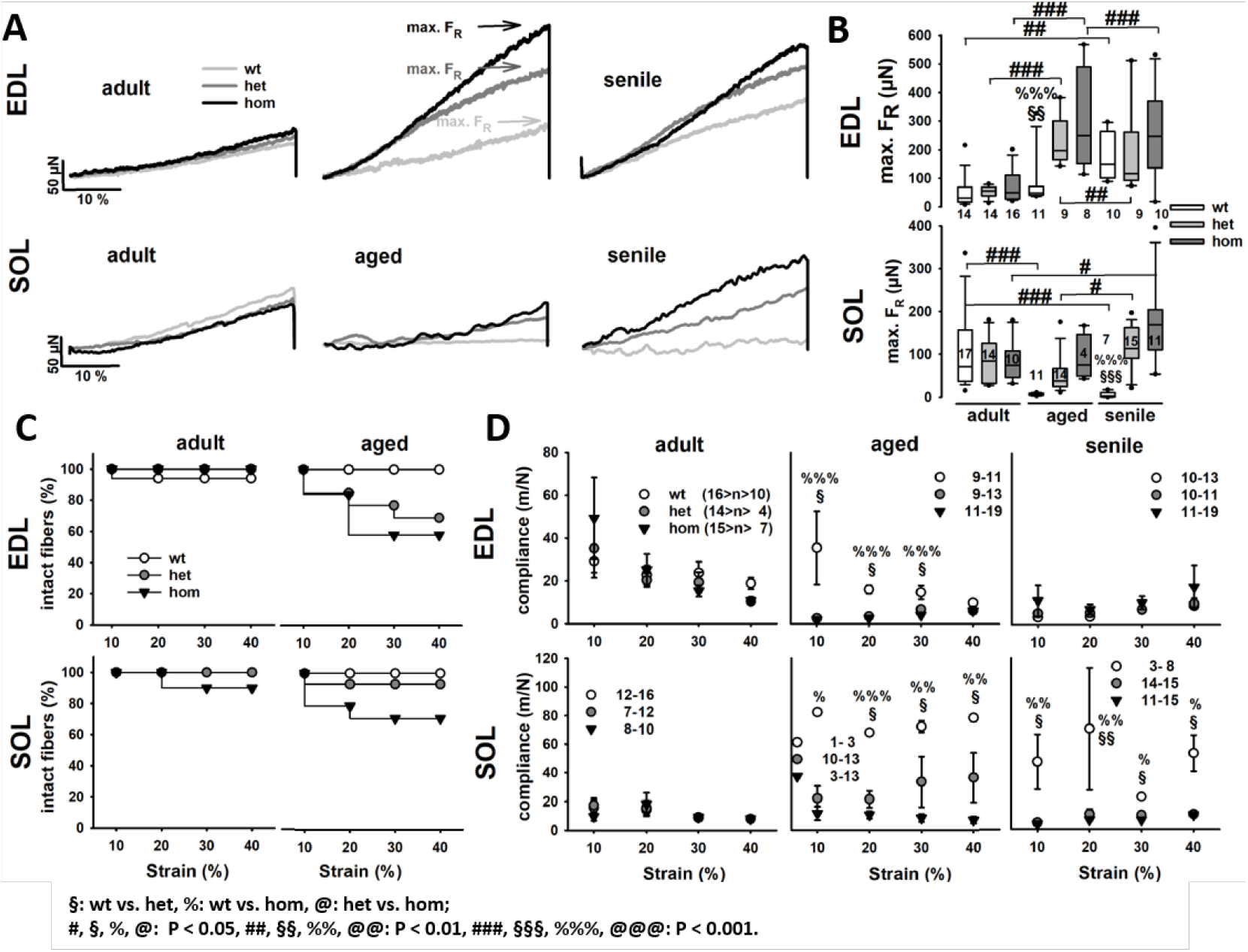
Passive steady-state resting length-tension curves in permeabilized single EDL and SOL fiber segments from adult, aged and senile mice carrying the R349P desmin mutation show a markedly increased axial mechanical stiffness and reduced compliance in mutant fibers. **A**, representative force recordings in single EDL (top) and SOL (bottom) fibers for each genotype and age bin investigated. During ageing, het and hom R349P desmin knock-in fibers present with a markedly steeper curve and increased maximum restoration forces FR. This was confirmed in the group analysis in (**B**), showing significantly increased FR values in both het and hom fibers already in the aged mice, while wt fibers remained reduced, but eventually increased within the senile age group. **C**, Kaplan-Maier survival plots, shown for the adult and aged group, depict a much lower survival of single fibers during the stretch protocol compared to wt fibers. **D**, mechanical axial compliance values derived from slopes to the RLT curves to 10% stretch bins show (i) marked decrease in compliance with stretch and (ii) much lower compliance values for mutant fibers over wt fibers except for adult mice in both EDL and SOL, and senile mice in the EDL muscle. Numbers in box plots: number of single fibers analysed.

### Axial viscosity is unaltered by the R349P mutation in single EDL and SOL fibers

While the R349P mutation clearly affects axial elasticity, RLT curves did not provide insights into the biomechanical axial viscosity. Therefore, we used the *MyoRobot* to perform ultra-fast stretch-jumps such as shown in Fig. 4A for wt adult single fibers from EDL and SOL muscle. Each new stretch jump was answered by an instantaneous restoration force (F_*R*_) increase to a maximum, followed by viscous relaxation (F_*relax*_) to a new steady-state during the 5 s holding phase. Confirming the findings from the slow RLT curves, mutant single fibers had a much higher chance of rupture during these strenuous sudden stretches as compared with wt fibers (Fig. 4B). The analysis of maximum F_*R*_ amplitudes with stretch bin, reflecting axial elasticity, confirmed the findings from the RLT curves, i.e. higher restoration forces in the mutant background (Fig. 4C). However, relaxation force F_*relax*_, representing the difference between maximum F_*R*_ and steady-state F_*R*_ within the same stretch jump, was not significantly different between either genotype or ages (Fig. 4D), arguing against any involvement of the R349P mutant desmin in viscous relaxation.

**Figure 4.**
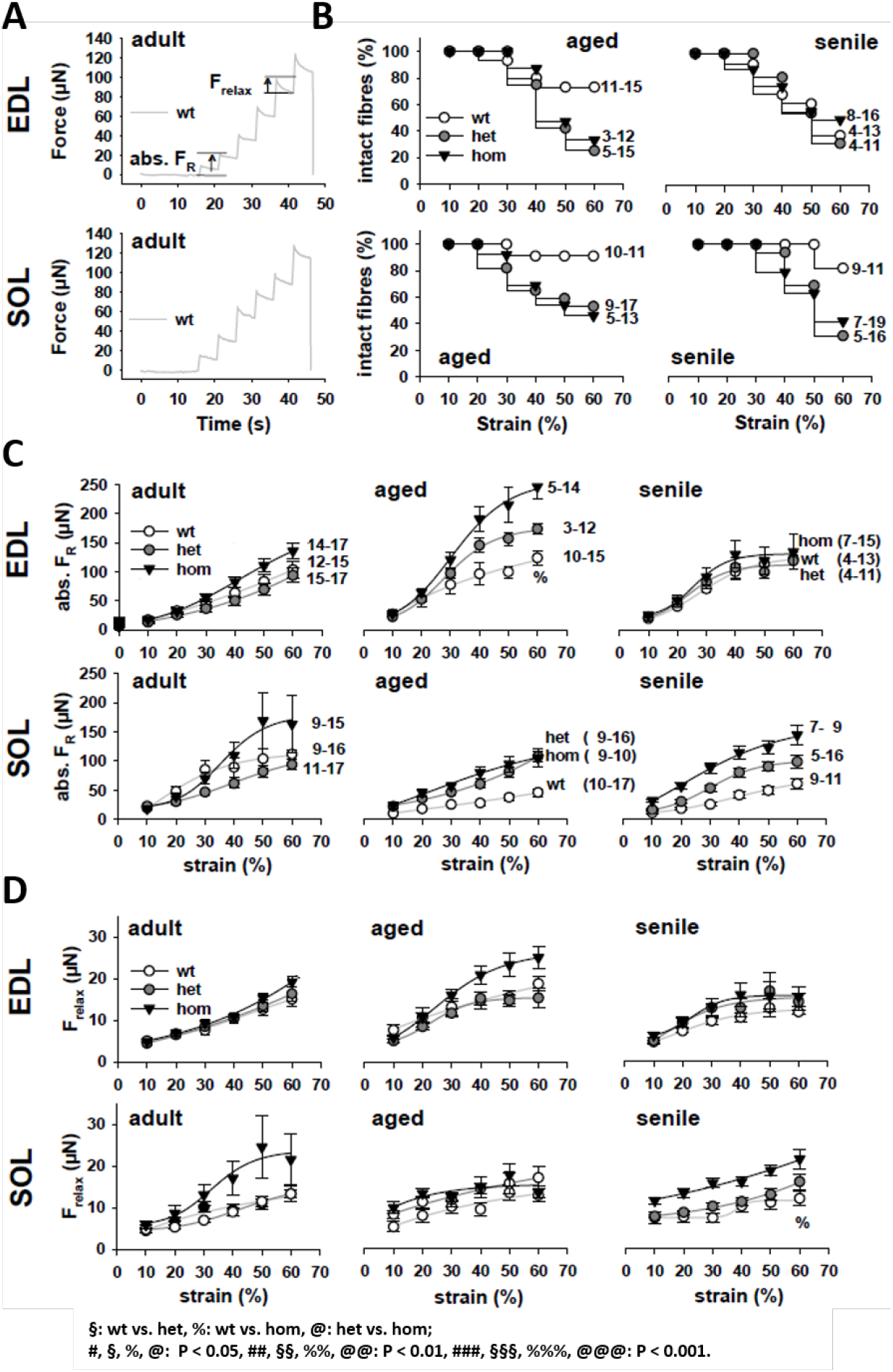
Visco-elastic behaviour of single muscle fibers from EDL and SOL muscle carrying the R349P desmin mutation during ageing. **A,** representative examples of quick step-stretch experiment protocols stretching adult EDL and SOL fibers in 10% bins to 160% L_0_. **B,** Kaplan-Meier curves displaying the percentage of intact fibers during the protocol demonstrate a worsened stretch-resistance of mutant fibers. **C,** group analysis of F*_R_* across ages in EDL (top) and SOL (bottom) fibers showing overall increased absolute restoration force levels in the mutants over the wt for almost all ages and in both muscles. **D,** force relaxation amplitudes with stretch suggest almost similar viscous relaxation with a tendency for higher viscous relaxation in mutant fibers over the wt. Error bars: standard error.

### Unloaded speed of shortening in R349P desmin single fibers is increased rather than compromised

The observed increased passive elasticity in R349P mutant desmin single fibers suggests a negative influence on muscle contraction kinetics, e.g. unloaded speed of shortening. To address this question, we performed so-called ‘slack-tests’ using our automated *MyoRobot* system ***Haug et al. (2018)***. Fig. 5A shows representative example recordings of a senile EDL (left) and an aged SOL (right) single fiber. After reaching steady-state maximum isometric contraction in HA solution, the VCA quickly introduced a slack of defined length dL. Consequently, force dropped to zero and redeveloped over time (dt). The relation dL vs. dt is plotted to the right in Fig. 5A. Also shown are the linearly derived fast and slow velocities v_*fast*_ and v_*slow*_ from the respective section of the double exponential fit. Fig. 5B shows the dL-dt plots for all age groups and genotypes for both muscles and Fig. 5C the statistical analysis of v_*fast*_ and v_*slow*_. V_*fast*_ reflects the initial, unloaded phase, whereas v_*slow*_ represents the internally loaded phase that occurs while taking up larger ‘slacks lengths’ ***Haug et al. (2018)***. Notably, v_*fast*_ increased with age in all genotypes, while it decreased again in senile mutation-bearing fibers, except for hom SOL fibers. In this context it was even more compelling that mutant fibers performed significantly faster than wt fibers in aged animals. Although v_*slow*_ qualitatively showed a similar trend, there were no statistical significances regarding age or genotype.

**Figure 5.**
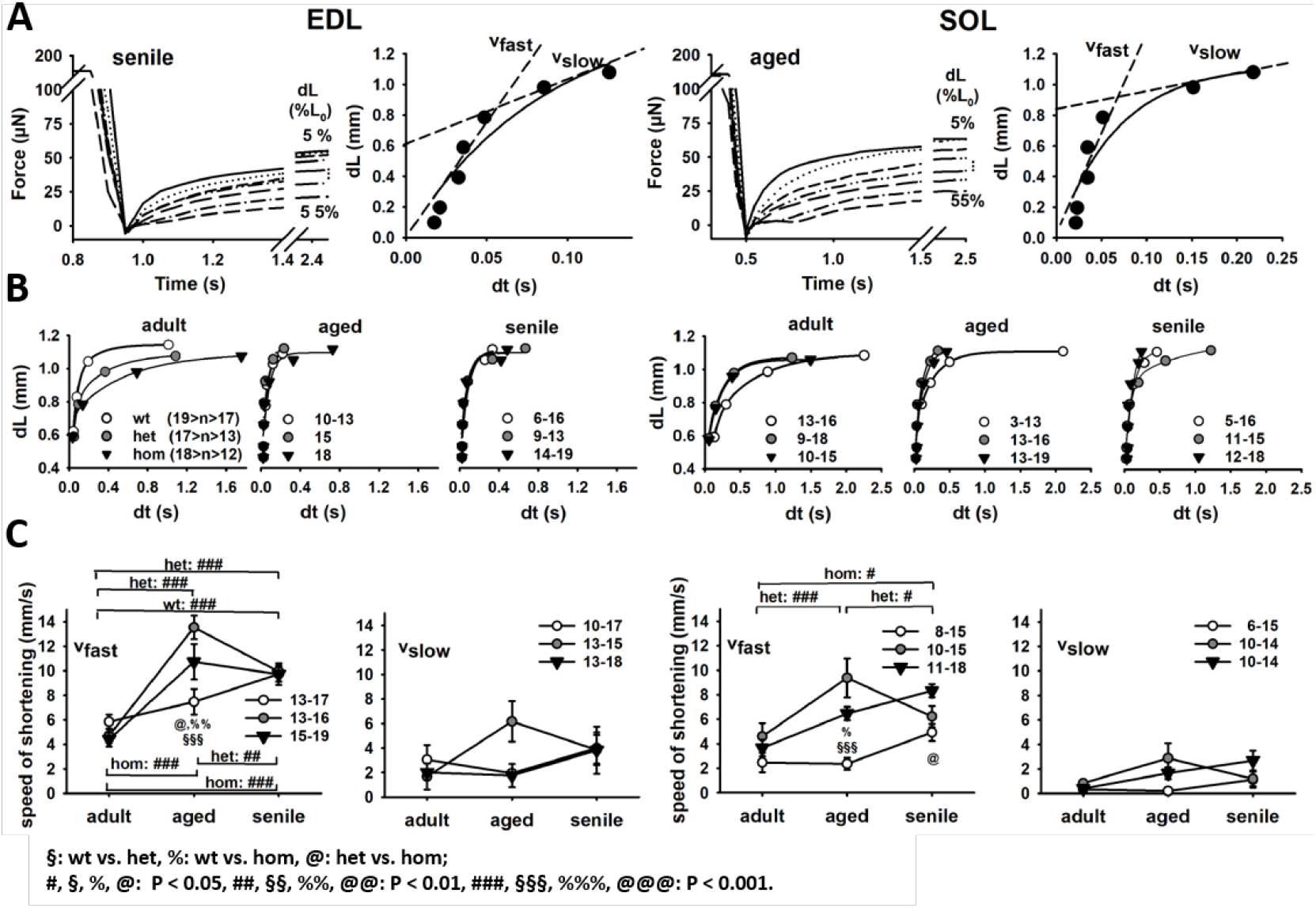
The fast phase of unloaded speed of shortening in single EDL and SOL fiber segments from adult, aged and senile R349P desmin mice is markedly increased in aged mutation-bearing fibers. **A,** representative ‘slack-test’ recordings of a single senile EDL (left) and aged SOL (right) fiber. The ‘slack time’ was extracted for each ‘slack length’ and the dL-dt relationship plotted in the right subpanels along with a biexponential fit and a linear velocity approximation in the lower dL (fast) and upper dL (slow) regime. **B,** group analyses of all single fibers from each genotype and age described by biexponential fit curves. The group analysis of the linear fast (v*_fast_*) and slow (v*_slow_*) phase for all fibers of each genotype and muscle is shown in (**C**). Shortening speed consistently increased in wt fibers with age, while particularly het samples reveal a maximum shortening velocity for aged fibers. In hom fibers, shortening speeds also increased with age and only displayed a single decline for senile EDL muscle. Numbers next to symbol legends: number of single fibers analysed.

## Discussion

Desminopathies comprise a heterogeneous group of inherited and sporadic myopathies which, in the vast majority of cases, share a common morphological picture comprising sarcoplasmic and subsarcolemmal desmin-positive protein aggregates and signs of myofibrillar degeneration ***JurcuŢ et al. (2017); Clemen et al. (2009); Walter et al. (2007)***. In general, analyses of the pathophysiology of human desminopathies are hampered by the very limited amount of available human muscle tissue specimens and the fact that the alterations noticed in diagnostic muscle biopsies nearly always reflect late stages of the disease. In addition, these specimens are highly heterogeneous with regard to sex, age, type of muscle and disease severity. To overcome these limitations, we generated the patient-mimicking R349P desmin knock-in desminopathy mouse model, which harbours the orthologous of the most frequent human desmin mutation R350P ***Clemen et al. (2015)***. The availability of this mouse line has already served invaluable to perform age-related morphometric analysis of cytoarchitectural changes in early disease stages in single fibers from slow- and fast-twitch muscles using multiphoton Second Harmonic Generation (SHG) microscopy ***Diermeier et al. (2017a)***. In that study, we could show a pre-aged morphological phenotype depicting sarcomeric lattice disorder and myofibrillar angular distribution in both EDL and SOL single fibers ***Diermeier et al. (2017a)***. On a single fiber level, such distorted myofibrillar cytoarchitecture would already be a structural determinant of muscle weakness ‘per se’ since the resulting force vector of the parallel myofibrillar lattice in single fibers would be expected to be smaller compared to if all myofibrils were perfectly aligned ***Friedrich et al. (2010); Buttgereit et al. (2013); Schneidereit et al. (submitted 2018)***. For human R350P desminopathy, apart from clinical assessment of overall force in proximal and distal muscle groups according to MRC grades ***Walter et al. (2007)***, no information on active force production on the sub-organ level (single fibers, fiber bundles) is available. For the murine R349P desmin knock-in model, initial characterization of small SOL fiber bundles at preclinical stages in young mice ***Clemen et al. (2015)*** as well as very recently, a whole age-dependent study of ours on small EDL and SOL fiber bundles in mice from 17wks to >60wks of age, documented a pre-aged phenotype regarding increased passive axial stiffness, using our novel high-end *MyoRobot* biomechatronics system ***Hauget al. (2018)***. Together with the finding of also increased extracellular fibrosis in aged R349P desmin knock-in muscles ***Diermeier et al. (2017a)***, this still leaves the possibility of series-elastic elements of increased stiffness in parallel to the cytoskeletal visco-elastic elements to be responsible for the reduced axial compliance in small EDL and SOL fiber bundles seen before. In order to close this gap, we refined our previous study to determine the active and passive biomechanics properties in het and hom R349P desmin knock-in mice as well as their wt littermates extending the age range from adult (35-45wks) to very old (90-96wks), as well as to mechanically dissected single fibers, for the very first time. The advantage of dissected single fibers not containing ECM connections to surrounding elements any more, i.e. being void of neighbouring fibers, provides a pure preparation to exclusively focus on the effect of mutated desmin on cytoskeletal axial fiber biomechanics.

### Age is the more predominant determinant of compromised active axial biomechanics in single fibers from slow- and fast-twitch muscle compared to the presence of R349P mutant desmin protein

The most important finding in our age-related study in single EDL and SOL fibers from adult to senile mice was that age had a strong negative influence on active force production, with significant declines during ageing within each genotype regarding the caffeine-induced Ca^2+^-mediated force transients. This effect was more prominent in slow SOL fibers but also, to a lesser extent, in fast-twitch EDL fibers. Among the genotypes, the presence of R349P mutated desmin was much less of a systematic determinant of caffeine-induced active force, in particular in SOL fibers, while in EDL fibers, some inconsistent significances were present. Detailed systematic age-related studies on contractile properties in fast- and slow-twitch muscle are rare with most studies having been carried out on whole muscle. In 2yr-old versus 6 mo-old rats, twitch and tetanic force were lower in old over young animals, but accounting for an age-related atrophy of fibers, no differences were found in the maximum force-generating capacity in either slow- or fast-twitch muscles at either age ***Larsson and Edström (1986)***. In the respective mouse muscles however, a decline in absolute isometric tetanic force production to ~75% of the values in young (2-3 mo) and adult (9-10 mo) was reported in aged (26 - 27 mo) mice for both EDL and SOL muscle. This difference prevailed after normalization to specific force for fast-twitch EDL, but for SOL specific tetanic force no more age dependence was seen ***Brooks and Faulkner (1988)***. In a comparative study on maximum isometric tension, comparing both fiber bundles and single skinned fibers from rat SOL and EDL muscle with age, maximum tension was increased by roughly 35% in senescent rats (30 mo) over control adults (9 mo) for both SOL bundles and single fibers. The same only applied for single EDL fibers whereas in EDL bundles, force almost dropped ~20% with age ***Eddinger et al. (1986)***. Lastly, in an age-related study in dystrophic mdx mice, single skinned SOL and EDL fibers displayed no differences comparing young (3-6wks)and adult (17 - 23 wks) animal ***Williams et al. (1993)***. The recognition of considerable variability in specific isometric force values between study groups has been stated to render comparisons between whole muscles, fiber bundles and single fibers with respect to ageing difficult ***Brooks and Faulkner (1988)***.

The important strength of our approach lies in the age-dependent assessment of R349P mutant desmin effects in an age-dependent background which was not available before. One limitation of our *MyoRobot* system at the time this study was initiated was still the lack of in-built optics to measure the fiber diameter for conversion of absolute force to specific, cross-sectional area (CSA)-normalized force. Towards the completion of data collection for the age-related biomechanics assessment here, ongoing optical engineering in our labs resulted in a more advanced version of the *MyoRobot* that now contains in-built optics and a CCD camera to capture fiber diameter online. Although this system will be presented elsewhere, the absolute single fiber force levels presented here are well in the range of those reported by Stelzer & Widrick (2003) ***Stelzer and Widrick (2003)*** in SOL fibers from adult (8-12wks) mice, around 150*μ*N per fiber at maximum Ca^2+^ activation. Inversely, when measured with our current setup, single mouse fibers from animals unrelated to this study, fiber diameter values of mostly between 30 *μ*m and 40 *μ*m were found which translates specific forces to roughly 15 N/cm^2^ or 150 kPa, in perfect agreement with single fiber specific force values from literature ***Williams et al. (1993); Stelzer and Widrick (2003)***.

Regarding the Ca^2+^ sensitivity of the contractile apparatus, a pre-aged phenotype in the R349P background was observed. Particularly mutation-bearing EDL fibers displayed a myofibrillar Ca^2+^-sensitization already within the aged age group, while in wt littermates, this became only apparent in the senile group. This corroborates well with results from our very recent age-related biomechanical assessment of R349P desmin small fiber bundles, where a very similar desensitization of about 0.2-0.3 pCa units was seen from the adult to the aged age group in EDL bundles, both the het and the hom ***Haug et al. (2019)***. For SOL, the single fiber data here do not seem to show a firm consistent trend among the genotypes with age, apart from a large scattering between individual SOL fibers. This could be due to marked differences of pCa_50_ values between fast- and slow-twitch fibers being present in the SOL muscle as it contains an almost equal proportions of either fiber type ***Edgerton et al. (1975); Lynch et al. (1993)***. Unlike in previous studies using single fiber Ca^2+^ sensitivity assessment (e.g. ***Lynch et al. (1993) Lynch et al. (1993)***), we did not attempt to type fibers for myosin heavy chain (MHC) isoforms due to technical reasons and thus, this may at least partially explain the observed scatter. However, from our previous work assessing the MHC composition in SOL muscle homogenates in the three genotypes, we are confident that in particular the hom fibers shall present with a higher slow-type MHC I proportion over wt and het fibers ***Diermeier et al. (2017a)***. Therefore, the large scatter towards higher pCa_50_ values in aged and senile single SOL fibers (Fig. 2B) is in good agreement with the fact of higher pCa_50_ values in type I over type II fibers ***Lynch et al. (1993)***. Also, our absolute pCa_50_ values presented here are in good agreement with the aforementioned study ***Lynch et al. (1993)***.

### Passive axial biomechanics is shifted towards a pre-aged stiffer phenotype in single fibers by R349P desmin

Similar to our previous assessment in small fiber bundles ***Clemen et al. (2015); Diermeier et al. (2017a); Haug et al. (2019)***, single fibers showed a marked increase in passive restoration forces in RLT experiments, which was at least three-times larger in aged animals with R349P background. Restoration forces were significantly increased and axial compliance accordingly decreased, already in single EDL fibers from aged R349P desmin knock-in animals compared to the equivalent observations found in wt animals in the senile group, only. This clearly points to a pre-aged phenotype in fast-twitch muscle in the mutant desmin background. Notably, the increase in restoration force (decrease in compliance) in EDL bundles already happened in young animals and was more pronounced in hom R349P desmin knock-in mice, albeit not statistically significant ***Haug et al. (2019)***. Also, from the adult age, the differences were blunted among genotypes in bundles in the EDL, arguing in favour of an important difference compared to single fibers. This can only be explained by the abolishment of ECM components in the pure single fibers blunting the effects of an additional parallel series elastic element contributing to the axial steady-state compliance. In our SOL single fibers, a similar increase in passive axial stiffness in het and more pronounced in hom R349P desmin knock-in single fibers was seen. This is also much more clear-cut here compared to the presentation in SOL bundles. In the latter, the increase in stiffness with age was mostly seen in hom bundles only, but those being highly statistically stiffer over het R349P and wt bundles in aged mice ***Haug et al. (2019)***. The direct comparison of bundles ***Haug et al. (2019)*** and single fibers now being available suggests that ECM components likely introduce an additional compliance to the bundles, e.g. through elastic fibers, that would reduce the overall stiffness. Alternatively, the over-proportionally larger cross-sectional area in bundles carrying the R349P mutation may be dissipating restoration forces between both, intracellular (i.e. mutated desmin) and extracellular non-contractile elements. Although an increase in ECM collagen was detected in our previous study in R349P SOL bundles that was larger in hom over het bundles ***Diermeier et al. (2017a)***, the involvement of other, more compliant ECM components to be increased in the R349P desmin knock-in background cannot be ruled out and deserves further investigation. Nevertheless, our *MyoRobot* approach was able to extend our previous knowledge on R349P axial muscle stiffness to single fibers and also including a larger age range extending to the senile stage, not only with regard to the R349P desmin knock-in background, but also in particular including ageing effects in normal muscle. For instance, in both our EDL and SOL single fiber preparations from wt mice, axial compliance increased from the adult to the aged age groups to then remain mostly stationary in the EDL, or they declined again in the SOL within the senile groups (Fig. 3D). This is in good agreement with a comparative study on *tibialis anterior* mouse muscle single fibers and small fiber bundles, where single fibers from old mice showed a tendency for reduced elasticity moduli (reflecting smaller stiffness, larger compliance values, respectively, kPa, n.s.) ***Wood et al. (2014)***. More strikingly, the researchers showed that intrinsic stiffness of ECM increased with age as indicated by larger Young moduli in fiber bundles over single fibers, and in particular, a two-fold increased bundle stiffness in old versus adult *tibialis anterior* fiber bundles ***Wood et al. (2014)***. The same behaviour of increased modulus (quadratic modulus, kP/*μ*m^2^) values was shown in EDL fiber bundles over single fibers from young (7-9wks) wt mice ***Meyer and Lieber (2011)***. When comparing our axial single fiber compliance values to the corresponding values in small fiber bundles given in our associated study in adult and aged mice (***Haug et al. (2019a) Haug et al. (2019)***, i.e. for SOL ~1-4m/N and for EDL bundles ~1-6m/N), one can see that our single fibers consistently show higher compliance values, reflecting higher stiffness in bundles over single fibers. Meyer & Lieber (2011) ***Meyer and Lieber (2011)*** also provide an elegant experimental explanation for the increased stiffness in fiber bundles over single fibers and grouped single fibers, in that the ECM contribution to non-linear bundle elasticity is set out by spreading the sarcomere length distribution of individual fibers within the bundle. This superposes different RLT-curves from single fibers in a bundle to a non-linear resultant elasticity behaviour. This is most probably due to different lateral and axial forces acting on adjacent single fibers through ECM-mediated focal adhesion connections, i.e. integrins ***Gershlak and Black (2015)***. It is of note that the absolute values for axial stiffness (compliance) in our study and those aforementioned ones cannot be directly compared, as different methods were employed, and our system during that time of experiments in single fibers could not yet assess single fiber CSA and sarcomere lengths. When focusing on the influence of mutant R349P on passive axial compliance/stiffness, our current results support those obtained in fiber bundles ***Haug et al. (2019)*** in showing significantly reduced axial compliance indicative of stiffer phenotype with the important difference that in the bundles, this effect seemed less systematic in the respective age groups in both EDL and SOL muscles. In single fibers, compliance was clearly at least two-fold reduced in the R349P background over wt animals, starting from the aged age group in both muscles and staying diminished in SOL muscles while alleviating in the EDL muscle from senile animals. Thus, revealing the higher stiffness in single fibers in a more robust way over the respective fiber bundle preparations ***Haug et al. (2019)*** again points to the crucial involvement of ECM components which introduce additional non-linear elasticities to the axial compliance that are difficult to predict in the R349P desmin knock-in background. The larger presence of fibrosis in muscle tissue from aged mice hom for the R349P desmin allele clearly points towards the presence of such non-linear elasticities over het and wt muscles ***Diermeier et al. (2017a)***. The larger stiffness in pure single fibers in the R349P desmin knock-in background is also confirmed by their lower resistance to stretch and thus, lower survival upon stretch, either quasi-static (Fig. 3C) or dynamic (Fig. 4B). Similar to our findings in bundles ***Haug et al. (2019)***, visco-elastic properties were not significantly altered in R349P desmin knock-in single fibers. It should be noted that apart from our detailed study here, nothing is known about the influence of intermediate-filament mutations on the visco-elastic properties of fully differentiated muscle fibers.

### Unloaded speed of shortening in ageing: faster contractions of R349P desmin knock-in single fibers

With our recent implementation of a VCA within the *MyoRobot* ***Haug et al. (2018)***, it was now possible to address whether the markedly increased axial stiffness in single fibers in the R349P desmin knock-in background would impact on unloaded shortening, given the fact that the isometric maximum force development was only marginally affected. The absolute values of velocities for the fast component with mean values between 4mm/s and 12mm/s for EDL single fibers and 2 - 6mm/s for SOL fibers are well in the range of velocities reported for single EDL fibers from wt mice unrelated to this study ***Haug et al. (2018)***. This demonstrates the robustness of our automated biomechatronics system to assess active biomechanical properties in single fibers across studies and organ scales. Similar to our previous study in small fiber bundles, fast velocities increased with age in het mice while in the wt, they remained stationary ***Haug et al. (2019)***. However, in the aforementioned mentioned study of ours, only few numbers of observations were available for EDL muscle bundles, which complicates a robust comparison. Rather, for SOL bundles with higher numbers of observations, fast velocities showed a tendency for slowed shortening in the R349P knock-in background in young animals that was mostly abrogated in the adult and aged age group, except for a tendency of a faster shortening in the mutants ***Haug et al. (2019)***. Thus, it was intriguing to find that both, fast and slow components of shortening in single muscle fibers increased with age, the more so for the R349P knock-in background over the wt for adult and aged mice for both EDL and SOL fibers, while values again merged to similar levels in senile mice. Age-related values for unloaded shortening in single fibers are scarce in the literature. A study on rat EDL single fibers found an unchanged maximum shortening velocity was in adult (9 mo) versus senescent (30 mo) animals, whereas SOL fibers from old rats ***Eddinger et al. (1986)*** were slower. In human *vastus lateralis* single skinned fibers, shortening velocities were reduced in type IIA fibers but not type I fibers in older man, while the opposite was found for women ***Krivickas et al. (2001)***. For murine muscles, a detailed sex and age-related study is not available, to our knowledge. The reason for the increased speed of shortening in R349P desminopathy muscle fibers cannot unambiguously be explained at current, in particular in view of our recent finding that slow-type MHC I isoforms were upregulated in R349P desmin knock-in muscles while fast-twitch MHC II isoforms were downregulated ***Diermeier et al. (2017a)***. Thus, the R349P mutated desmin must have some influence on the kinetics of weak cross-bridge attachment that has been found to predominantly determine unloaded speed of shortening ***Stehle, R. & Brenner, B. (2000)***. Whether this may be an explanation for the increased shortening velocity in desminopathy single fibers deserves future investigation.

### Summary and outlook

Our results prove the increased passive steady-state elasticity found in R349P desminopathy skeletal muscle to be an inherent factor related to the mutant desmin inflicted damage of the cytoskeleton and not the ECM. This results in a pre-aged, stiffer phenotype of affected muscle fibers. Apart from a yet unexplained acceleration of speed of shortening in fibers expressing R349P desmin, Ca^2+^-mediated active force was only mildly affected, if at all. Our *MyoRobot* system allows a highly versatile and modular design of automated execution of various additional muscle test protocols, e.g. eccentric contractions, that shall be of use to the community to ease future myopathy and mechanistic studies related to skeletal muscle and ageing.

## Methods and Materials

### Mouse model - R349P Desmin knock-in mouse

Heterozygous (het) and homozygous (hom) littermates of the R349P desmin knock-in mouse model B6J.129Sv-Des^*tm*1.1*Ccra*^ (http://www.informatics.jax.org/allele/MGI:5708562) ***Clemen et al. (2015); Winter et al. (2016)*** were used. Littermates not carrying the R349P desmin mutation served as wild-type (wt) control. We here extended our previous biomechanics study on small fiber bundles from only young mutant mice ***Diermeier et al. (2017a)*** towards three older age groups, spanning 35-45 weeks (adult), 65-75 weeks (aged) and 90-96 weeks (senile). All animal-related work was performed in accordance with the German Animal Welfare Act (Tierschutzgesetz), as well as the German Regulations for the protection of animals used for experimental purposes or other scientific purposes (Tierschutz-Versuchstierverordnung). The governmental Office for Animal Care and Use (Regierung von Mittelfranken, 91511 Ansbach, Germany; reference number TS-14/2015) approved the investigations. All applicable international, national, and institutional guidelines for the care and use of animals were followed.

### Chemical solutions

All muscle dissection was performed in Krebs-solution containing (mM): 120 NaCl, 4.7 KCl, 1.2 KH_2_PO_4_,1.2 MgSO_4_x7H_2_O, 24.8 NaHCO_3_, 0.1 M glucose, 0.1% FCS (FBS), pH 7.3. A Ca^2+^-free, high K^+^-solution (HKS) was used to permanently depolarize the muscle cell membrane to abolish excitability during manual tethering of fascicles and isolation of single fiber segments. HKS (‘high-K^+^-solution’) contained 140 K-glutamate, 10 Hepes, 10 glucose, 10 MgCl_2_,1 EGTA(ethylene glycol-bis(*β*-aminoethyl ether)-N,N,N’,N’-tetraacetic acid, pH 7.0. To maximally Ca^2+^-activate single fibers, a Ca^2+^-saturated high activating internal solution (HA) was used containing: 30 Hepes, 6.05 Mg(OH)_2_, 30 EGTA, 29 CaCO_3_, 8 Na_2_ATP, 10 Na_2_CP, pH 7.2. Free Ca^2+^ of HA was calculated to ~12.5 *μ*M using the chelator-ligand binding software React (developed by Geoffrey Lee, University of Glasgow). To maximally relax single fibers and to completely buffer Ca^2+^ ions each time a fiber was exposed to Ca^2+^, high relaxing solution (HR) was used that had the same composition as HA except for not containing any Ca^2+^ (for practical reasons of pCa calculations, a pCa of 9 is assumed in HR). Mixtures of HA and HR were calculated to obtain a given pCa of the internal solution for graded Ca^2+^-activation in pCa-force response curves using React and consisted of HA:HR ratios of 0.3:0.7, 0.5:0.5, 0.55:0.45, 0.6:0.4, 0.65:0.35, 0.7:0.3, 0.8:0.2, 0.9:0.1, 0.95:0.05, 0.98:0.02, 1:0, converting to pCa values of 6.74, 6.38, 6.30, 6.21, 6.12, 6.03, 5.82, 5.54, 5.32, 5.11, 4.92, respectively. Low relaxing solution (LR) served as an intermediate step after HR or loading solution (LS, see below) to replace the high affinity Ca^2+^ chelator EGTA for low affinity HDTA (1,6-diaminohexane-N,N,N’,N’-tetraacetic acid). LR contained: 30 Hepes, 7.86 Mg(OH)_2_, 87.8 K-glutamate, 6.6 HDTA, 0.4 EGTA, 8 Na_2_ATP, 10 Na_2_CP (creatine phosphate), pH 7.2. LS was a mixture of HA and HR titrated to a free Ca^2+^ of ~300 nM to reload the sarcoplasmic reticulum (SR) for defined incubation times. RS served as release solution for Ca^2+^ ions from the SR and was LR supplemented with 30 mM caffeine. All solutions were thawed from stocks at the day of experiments and freshly supplemented with creatine kinase (CK, Sigma/Roche, Germany) to ~300 U/ml or ~3 U/well and sodium azide (0.1 M NaN_3_), the latter to prevent mitochondrial Ca^2+^ uptake ***Fry et al. (1989)***. To initially chemically permeabilize a single fiber, saponin was added to HR in a separate well of the *MyoRobot* rack to a concentration of 0.1% (w/v).

### Preparation of single muscle fibers

Mice were anaesthetized via isoflurane inhalation and sacrificed by cervical dislocation. The hind limbs were cut off and transferred to Krebs-solution. SOL and EDL muscles were dissected under a stereo-microscope (Olympus SZX7, Olympus, Hamburg, Germany), while being pinned under slight stretch into a Sylgard (Dow Corning, Wiesbaden, Germany) coated petri dish. Upon completing the dissection, Krebs-Solution was exchanged for HKS, allowing for 15 min equilibration, before single fibers were manually dissected with fine forceps.

### Assessment of active and passive biomechanics in single muscle fibers in an automated *MyoRobot* environment

Biomechanics recordings were conducted using the *MyoRobot*, a novel automated biomechatronics system combining high precision voice coil actuation (VCA) with force sensor technology ***Haug et al. (2018)***. After isolation, the single fiber segment (length at least 2 mm) was transferred to the *MyoRobot* multi-well rack in a custom-made Perspex chamber while submerged in HKS solution, placed below the pins of the force transducer and VCA and fixed to both pins via a tweezer mechanism. For details on the biomechatronics system and sensor, and actuation implementation, refer to ***Haug et al. (2018)***. Every protocol started with a chemical permeabilization of single fibers in HR supplemented with saponin for 20 s. An automated set of biomechanical recordings on the same preparation was then executed, consisting of sequential runs of (i) caffeine-induced, Ca^2+^-mediated force generation, (ii) pCa-force curves, (iii) speed of shortening (slack-test), (iv) passive elasticity - resting length-tension curve (RLT) and (v) assessment of visco-elastic passive behaviour:

- **Caffeine-induced, Ca^2+^-mediated force generation:** After fiber permeabilization, the fiber was shortly dipped into HR to wash off remaining saponin and to buffer internal Ca^2+^. Subsequently, it was translocated to LR for 60 s, after which the SR was loaded in LS for 60 s. The caffeine-induced force transient was triggered by exposure to RS for 60s, while maximum force was induced via HA solution for 5 s (Fig. 1).
- **Ca^2+^ sensitivity of the contractile apparatus, pCa-force curves:** The fiber was sequentially exposed to solutions of increasing Ca^2+^ ion concentrations (decreasing pCa values (−log_10_[Ca^2+^])) for a duration of 20 s (Fig. 2).
- **Unloaded speed of shortening (‘slack-test’):** The muscle fiber was held at resting length L_0_ and transferred to HA solution, resulting in maximum isometric contraction. Upon achieving steady-state force, the VCA pin moved at maximum speed towards the force transducer, slacking the fiber by a defined percentage of L_0_ (5%, 10%, 20%, 30%, 40%, 50% or 55%) as force dropped to 0mN. While taking up the slack, force re-established in the presence of saturating Ca^2+^. Once the next force-plateau was reached, the fiber was washed in HR to remove excessive Ca^2+^ and to relax the myofibrils before moving on to the consecutive ‘slack length’. For this recording, sampling rate was set to 2 kHz (Fig. 3).
- **Passive elasticity - RLT curves:** To assess passive axial elasticity, the muscle fiber was kept in LR solution to avoid active contraction. The fiber was continuously stretched at a slow speed (0.44 *μ*m/s) to 140% of L_0_ (L_0_ ~1,950*μ*m) by moving the actuator pin away from the force transducer pin. Restoration force was continuously recorded. To every 10% stretch bin, a linear fit was applied to calculate the fiber’s compliance, reflected by the inverse of that slope and thus, the inverse of stiffness (Fig. 4).
- **Visco-elastic passive behaviour:** To assess the visco-elastic passive behaviour, the fiber was stretched in a sudden staircase-like pattern in 10% L_0_ steps to 160% L_0_ with a holding time of 10 s. To prevent any active contraction, the fiber was kept in LR during the recording. The force response of the fiber comprised of an instantaneous passive restoration force and a force relaxation, with an exponential decay of force back to a steady-state level (Fig. 5).

### Data analysis and statistics

*MyoRobot* data were processed with analysis protocols in RStudio (RStudio Inc., rstudio.com) while plotted and statistically evaluated with SigmaPlot (Systat Software Inc., sigmaplot.co.uk). All data traces were smoothed with a moving average filter. For pCa-curves, the plateau force close to the end of each pCa step was determined by the software and plotted against the corresponding pCa value. The scatter plot of normalized force (normalized to max. force at pCa 4.92) was fitted to a four-parameter Hill-equation 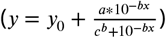 utilizing least-square methods with the physiological constraints y_0_ = 0 and a = 1. The steepness (b, Hill-coefficient) and the deflection point (−*log*_10_([*Ca*^2+^]), pCa_50_) of every individual curve fit were used to reconstruct a mean fit to the averaged data points (Fig. 2). For speed of shortening (slack-tests), a 5% threshold criterion was established from the maximum isometric force of the first slack. This threshold defined significant ‘force redevelopment’ for this and all consecutive ‘slack lengths’ (dL). The time needed to cross this force-threshold was called ‘slack time’ (dt) and was plotted against the respective ‘slack length’ dL. The resulting dt-dL-scatter plot was fitted with a bi-exponential function (*y* = *a*(1 − *e*^*κ*1+*dt*^) + *c*(1 − *e*^*κ*2+*dt*^)). Its derivative represented the non-linear slack-length dependent shortening velocity v(dL). The dL-dt range was divided in a fast (unloaded phase, <45% ‘slack length’) and a slow phase (internally loaded phase, >45% ‘slack length’) as described in Haug et al. (2018) ***Haug et al. (2018)***. Passive elasticity - RLT curves: To every 10% L_0_ stretch bin, a linear ft was applied and the respective increase/steepness computed to obtain axial elasticity and compliance (inverse increase). Viscoelastic behaviour: The force baseline (F_0_) was determined as the last 5 s before the first step while absolute restoration force (*F_abs_* = *max*_*n*\*10%_ − *F*_0_) of each 10% stretch-step was calculated as the difference of maximum recorded force of the corresponding step to the baseline. Force relaxation was obtained from the difference between maximum and minimum force recorded within the same step (*F_relax_* = *max*_*n*\*10%_ − *min*_*n*\*10%_). Statistical significance was assessed by applying two-way ANOVA tests (age bins and genotypes as variables), following post-hoc analysis (Tukey, Bonferroni) in SigmaPlot. Significance levels of P < 0.05 were considered significant, < 0.01 and < 0.001 considered strongly and highly significant, respectively. Significance levels involving age effects were depicted as #, while genotype differences were depicted as §: wt vs. het, %: wt vs. hom, and @: het vs. hom, respectively.

## Acknowledgments

This study was supported by a project grant from the Deutsche Forschungsgemeinschaft FOR1228 (to O.F., R.S., C.S.C.), a grant from the Central Innovation Program SME of the German Ministry of Economy & Technology (ZIM-Kooperationsprojekt KF2347924AK4) to O.F. and by funds through the Staedtler-Stiftung DS/eh 11/14 and Marohn-Stiftung MyoDes to OF, RS.

## Conflict of interest

The authors declare that they have no competing or financial interests.

## Author contributions statement

C.M., M.H. and B.R. conducted the experiments. C.M. analyzed the results. G.P. and M.H. engineered the *MyoRobot* biomechatronics system. R.S. and C.S.C. provided the animal model and expertise on the pathophysiology of desminopathiess, and O.F. conceived the project and supervised the whole research. C.M., M.H., R.S., C.S.C and O.F. wrote the manuscript. All authors approved the manuscript. This paper is part of the doctoral thesis of C.M.

## Additional information

O.F. discloses relationship to the SME conmoto through the aforementioned ZIM grant as an R&D project to translate the *MyoRobot* technology into commercialization.

